# The influence of heart rate variability biofeedback on cardiac regulation and functional brain connectivity

**DOI:** 10.1101/2020.08.10.243998

**Authors:** Andy Schumann, Feliberto de la Cruz, Stefanie Köhler, Lisa Brotte, Karl-Jürgen Bär

## Abstract

**Background:** Heart rate variability (HRV) biofeedback has a beneficial impact on perceived stress and emotion regulation. However, its impact on brain function is still unclear. In this study, we aimed to investigate the effect of an 8-week HRV-biofeedback intervention on functional brain connectivity in healthy subjects.

**Methods:** HRV biofeedback was carried out in five sessions per week, including four at home and one in our lab. A control group played *jump‘n’run games* instead of the training. Functional magnetic resonance imaging was conducted before and after the intervention in both groups. To compute resting state functional connectivity (RSFC), we defined regions of interest in the ventral medial prefrontal cortex (VMPFC) and a total of 260 independent anatomical regions for network-based analysis. Changes of RSFC of the VMPFC to other brain regions were compared between groups. Temporal changes of HRV during the resting state recording were correlated to dynamic functional connectivity of the VMPFC.

**Results:** First, we corroborated the role of the VMPFC in cardiac autonomic regulation. We found that temporal changes of HRV were correlated to dynamic changes of prefrontal connectivity, especially to the middle cingulate cortex, left anterior insula, right amygdala, supplementary motor area, dorsal and ventral lateral prefrontal regions. The biofeedback group showed a drop in heart rate by 5.5 beats/min and an increased RMSSD as a measure of HRV by 10.1ms (33%) after the intervention. Functional connectivity of the VMPFC increased mainly to the right anterior insula, the dorsal anterior cingulate cortex and the dorsolateral prefrontal cortex after biofeedback intervention when compared to changes in the control group. Network-based statistic showed that biofeedback had an influence on a broad functional network of brain regions.

**Conclusion:** Our results show that increased vagal modulation induced by HRV-biofeedback is accompanied by changes in functional brain connectivity during resting state.

## Introduction

The heart is the central organ of the circulatory system that pumps blood through the arterial vessel network in order to provide oxygen for all vital organs. Although the activity of the heart is driven by an intrinsic pacemaker called sinoatrial node, it is additionally influenced by environmental demands. Body signals shape the workload of the heart in order to meet changing needs of the entire organism. The two peripheral branches of the autonomic nervous system (ANS), the parasympathetic and the sympathetic system, modulate the intrinsic activity of the cardiac pacemaker cells in the sinoatrial node. While the sympathetic branch is needed for an adequate stress response, parasympathetic or vagal activation reduces expenditure and promotes health. Thus, the heart rate mirrors the resulting homeostasis of an organism influenced by internal and external demands. It is, therefore, conceivable that a complex system is needed to orchestrate autonomic cardiac function.

On the basis of early animal experiments and lesion studies, Benarroch described the “central autonomic network” (CAN) including several regions of the forebrain, limbic system, and the brainstem[1]. In a previous meta-analysis, we found that core nodes of the CAN have been consistently reported by modern neuroimaging studies on regulation of the ANS, i.e. the cingulate cortex, anterior insula, ventromedial prefrontal cortex (VMPFC), mediodorsal thalamus, amygdala, hypothalamus etc[2]. Further reviews corroborate a cortico-limbic network, including VMPFC, cingulate cortex, insula, and amygdala, to drive central autonomic control[3,4].

As a more theoretical framework, Thayer and colleagues introduced the neurovisceral integration model that links cognitive and emotional states to autonomic function[5]. In their approach, the prefrontal cortex is the highest level of a hierarchical model with direct functional connections to limbic regions, i.e. the insula and cingulate[4,5]. The limbic system is further connected via the amygdala to subcortical downstream regions such as hypothalamus and brainstem nuclei that determine parasympathetic and sympathetic modulation of the heart, at the lowest level of the model. According to this construct, the prefrontal cortex and its top-down control over subcortical structures has a pivotal role in heart rate regulation and relates sympathovagal balance to cognitive and emotional processing[6].

Using magnetic resonance imaging and resting-state functional connectivity (RSFC) studies have corroborated the important role of the interaction between the medial prefrontal cortex and limbic regions in heart rate regulation[7,8]. In a recent publication, we compared RSFC patterns between groups of healthy individuals that differed in heart rate regulation. Our results indicated that subjects with slow heart rates have significantly increased RSFC in a functional network of several regions of the central autonomic and the sensorimotor system when compared to subjects with fast heart rate[9]. Interestingly, we observed an increased RSFC between the prefrontal cortex (VMPFC) and the anterior insula to be associated with slow heart rates.

Whereas heart rate is modulated by both branches of the autonomic nervous system, heart rate variability (HRV) is a marker of parasympathetic cardiac regulation. It is generally accepted that high variability of the heart rate is in many aspects health-promoting. Thus, lower levels of HRV have been associated with increased cardiovascular morbidity and mortality[10]. Hillebrand et al. (2013) reported that healthy subjects with diminished resting HRV have a 32–45% increased risk to suffer from a first cardiovascular event[11]. Furthermore, HRV is thought to be associated with cognitive performance and emotional well-being (see reviews[12,13]). Whether these correlational associations describe HRV as a consequence of central regulatory processes or as a prerequisite for effective regulation is still unclear[14]. In a recent opinion paper, Thayer and Mather (2018) proposed that oscillations in emotion-regulating networks can be enhanced by HRV biofeedback[14]. HRV biofeedback is a bio-behavioral intervention to augment vagal tone. It is based on the phenomenon of respiratory sinus arrhythmia that describes heart rate to increase during inhalation and to decrease during exhalation. Thus, subjects can modulate their heart rate and thereby their HRV by modifying their breathing pattern.

Several studies demonstrated a beneficial effect of biofeedback in psychiatric disorders such as depression[15]. A meta-analytic review of 24 studies, including 484 participants showed that HRV biofeedback reduces perceived stress and anxiety levels significantly[16]. Thus, it has been speculated that HRV biofeedback influences brain function. Its similarity to electrical vagal nerve stimulation has been focused in order to explain underlying physiological mechanisms[17]. As a method of treatment in patients with epilepsy or depression, a pulse generator that is implanted in the chest wall stimulates primarily afferent vagal fibers. Impulses reach the brainstem and influence areas in the forebrain that are involved in the regulation of emotions, cognitive and autonomic function such including the frontal cortex, amygdala, and insula[17–19].

In this study, we aimed to investigate the effect of HRV biofeedback on functional brain organization. We hypothesized that HRV biofeedback slows resting heart rate and increases its variability. As we assume that HRV is closely tied to fronto-limbic connectivity, we hypothesized increased functional connectivity between the VMPFC and core regions of the limbic system, especially the anterior insula and the cingulate cortex, after biofeedback intervention.

## Methods

### Study group formation

A total of 24 healthy participants were recruited from the local community via flyers and online advertisement and assigned randomly to two different treatment groups. Fourteen participants performed a biofeedback training (7 males; 7 females; age: 30 ± 9 y, 22 – 52 y). Ten participants completed a control intervention (5 males; 5 females; age: 30 ± 13 y, 18 – 55 y). Pregnancy, the intensive pursuit of endurance sports, cardiovascular diseases (e.g., hypertension, diabetes), neurological disorders (e.g., migraine, epilepsy, multiple sclerosis), or psychiatric disorders (e.g., depression, attention deficit hyperactivity disorder, anxiety disorder) were held as exclusion criteria. All participants gave written informed consent to a protocol approved by the Ethics Committee of the medical faculty of the Friedrich-Schiller University Jena (# 5423-01/18) in accordance with the Declaration of Helsinki.

### Intervention protocol

The intervention took eight weeks in which participants of the biofeedback group performed an HRV biofeedback training. Five training sessions per week had to be conducted, including four sessions at home and one session at the laboratory. Subjects in the control group played one of three different *jump’n’run* mobile games in sessions organized according to the same schedule as the biofeedback group.

Heart rate was recorded using a sensor incorporated in a belt that was tied around the chest of the subject (H10/H7 Heart Rate Sensor; Polar Electro Oy, Kempele, Finland). Via Bluetooth, the application EliteHRV (Elite HRV LLC, 2017) collected data from the sensor, stored recordings and displayed heart rate. Participants in the control group recorded heart rate in the background while playing a mobile game. In the biofeedback group, heart rate oscillations were displayed on the screen of their smartphone as instantaneous visual feedback of heart rate. Participants were asked to adapt their breathing patterns in such a way to enhance heart rate oscillations, as described below. After each training session, we received raw data acquired during that session per email from participants of both groups. Thus, we were able to track the progress of the training throughout the intervention.

Magnetic resonance imaging (MRI) was conducted before the beginning of the training (T1) and after finishing the schedule (T2). One week prior to T1, an additional MRI session was planned to obtain participants’ habituation to the procedure (T0). Physiological signals and resting state scans were acquired simultaneously in order to assess cardiac autonomic function.

### HRV Biofeedback

In the biofeedback group, participants’ current heart rate (HR) was shown as an interpolated smoothed curve on their smartphone display. Participants were briefed to observe the curve as it changed with their breathing rhythm. As their final goal, participants were instructed to breathe ‘in phase’ with their HR curve by inhaling when HR ascended and exhaling when it descended in order to expand the amplitudes of the HR curve. For further details of the intervention that was designed following the manual published by Lehrer et al. (2000) we refer to our previous publication[21].

### MRI data acquisition

The data were collected on a 3T whole-body system equipped with a 12-element head matrix coil (MAGNETOM Prisma, Siemens Healthineers, Erlangen, Germany). Participants were instructed to keep their eyes open during the whole measurement and to move as little as possible. T2*-weighted images were obtained using a multiband multislice GE-EPI sequence (TR = 484 ms, TE = 30 ms, FA = 90°, Multiband Factor = 8) with 56 contiguous transverse slices of 2.5 mm thickness covering the entire brain and including the lower brainstem. The matrix size was 78 × 78 pixels with an in-plane resolution of 2.5 × 2.5 mm2. A series of 1900 whole-brain volume sets were acquired in one session. High-resolution anatomical T1-weighted volume scans (MPRAGE) were obtained in a sagittal slice orientation (TR = 2300 ms, TE = 3.03 ms, TI = 900 ms, FA = 9°, acquisition matrix = 256 × 256 × 192, acceleration factor PAT = 2) with an isotropic resolution of 1 mm^3^.

### Physiological recordings and analyses

During functional MRI data acquisition at rest, respiratory and cardiac signals were recorded simultaneously using an MR-compatible BIOPAC MP150 polygraph (BIOPAC Systems Inc., Goleta, CA, USA) and digitized at 500 Hz. Respiratory activity was assessed by a strain gauge transducer incorporated in a belt tied around the chest, approximately at the level of the processus xiphoideus. An optical finger pulse sensor was attached to the proximal phalanx of the index finger of the subject’s left hand. Inter-beat-interval (IBI) time series were derived from the pulse signal. Finally, subject’s mean HR was computed, as well as global (standard deviation of heartbeat intervals, SDNN) and short term (root mean square of successive heartbeat interval differences, RMSSD) measures of HRV[22]. Breathing rate (BR) was estimated as inverse of the average interval between respiratory maxima that were derived from the respiration signal. The quality of heartbeat and respiratory peak detection was visually inspected for artifacts and corrected manually when necessary.

### Resting state functional MRI preprocessing

Data preprocessing was performed using AFNI (https://afni.nimh.nih.gov/) and SPM12 (http://www.fil.ion.ucl.ac.uk/spm). The first twenty images were discarded, allowing magnetization to reach a steady state. Physiological noise correction was performed by including four low-order Fourier time series to reduce artifacts synchronized with the respiratory cycle[23] and five respiration volumes per time (RVT) regressors that model slow blood oxygenation level fluctuations[24,25]. The RVT regressors consisted of the RVT function and four delayed terms at 5, 10, 15, and 20 s[24].

Further preprocessing included realignment to the first volume using a rigid body transformation. For each participant, head movement was below 3 mm and 3°. Additional preprocessing steps were (i) removal of lineal and quadratic trends and of several sources of variance, i.e. head-motion parameter, CSF and white matter signal, (ii) temporal band-pass filtering, retaining frequencies in the 0.01-0.1 Hz band, and (iii) spatial smoothing using a Gaussian kernel of full-width half maximum of 6mm. Extra-cerebral tissue was removed from the anatomical images using ROBEX[26], a learning-based brain extraction method trained on manually “skull-stripped” data from healthy subjects. These skull-stripped brains were aligned to the standard MNI 2-mm brain. Finally, functional images were registered to anatomical data and normalized to the MNI space by applying transformation parameters derived from the anatomical to MNI registration.

### Region of interest and functional connectivity analyses

Based on our hypothesis, a region of interest was defined in the VMPFC as the seed region for functional connectivity analyses. The VMPFC-ROI was drawn as a sphere of 10mm radio centered at MNI-coordinates, x=0, y=44, z=−14, as defined in our previous study[9].

To obtain functional connectivity maps, preprocessed resting-state fMRI signal was averaged over each voxel with the seed region and correlated against all voxels in the brain. The resulting Pearson correlation coefficients were converted to Fisher z statistics in order to produce a more normally distributed variable[9].

The effect of biofeedback training was evaluated comparing VMPFC correlation maps at T1 to T2 (paired t-Test). The effect of group (biofeedback vs. control) on RSFC changes between T1 and T2 was assessed using a two-sample t-test of z-map differences (T2-T1). Statistical results were thresholded with p<0.005 uncorrected at voxel-level and cluster size k>10 voxels[27].

### Sliding-window analysis of functional connectivity and heart rate variability

The correlation of functional connectivity changes with temporal changes of HRV was estimated at T1 using data from all participants (N=24). We calculated short-term variability of heart rate (RMSSD) and functional connectivity of the VMPFC ROI in sliding time windows of 45s length (90 TR) with 50% overlap (45TR)[28]. On the subject level, we performed a linear regression analysis of VMPFC RSFC maps and the HRV regressor. Contrast images were then passed into a one-sample t-test group analysis. The statistical map was thresholded at an uncorrected voxel-level significance of p < 0.005 and cluster size k>10 voxels.

### Network analysis of Resting state functional MRI

In addition to the seed-based FC approach, we investigated significant between-group differences in the whole-brain network connectivity (connectome) using the network-based statistic approach (NBS)[29].

Individual connectivity matrices were generated extracting the mean time series from 260 independent anatomical ROIs, which were defined based on the coordinates from an extensively validated parcellation system provided by Power et al.[30]. Each ROI was modeled as 10 mm diameter sphere with a minimum distance of 10 mm between sphere centers, thus avoiding potential overlapping. In addition, we discarded short-distance correlations less than 20 mm since it might be affected by spatial smoothing or reslicing. A paired t-test design was then performed on each group separately by comparing T2 vs. T1. Here, components were identified using a primary component-forming threshold at t > 4.6. Permutation testing (10 000 permutations) was then applied to calculate FWE for every component previously identified. Results were considered significant for *p* < 0.05. NBS analysis was conducted using the Brain Connectivity Toolbox[31].

## Results

### Temporal co-variation of prefrontal connectivity and heart rate variability

Pooling data from all participants (n=24) prior to the intervention (T1), we aimed to corroborate the association of heart rate variability and connectivity of the prefrontal cortex irrespective of biofeedback. In sliding windows, synchronous changes of functional connectivity and RMSSD were estimated. Individual z-maps of the correlation between those time series were tested for a significant temporal co-variation (one-sample t-test). Dynamic changes of short-term HRV were correlated to changes of prefrontal connectivity, especially to the middle cingulate cortex, left anterior insula, right amygdala, supplementary motor area, dorsal and ventral lateral prefrontal regions (see Figure 1, Table S1).

**Figure 1.**
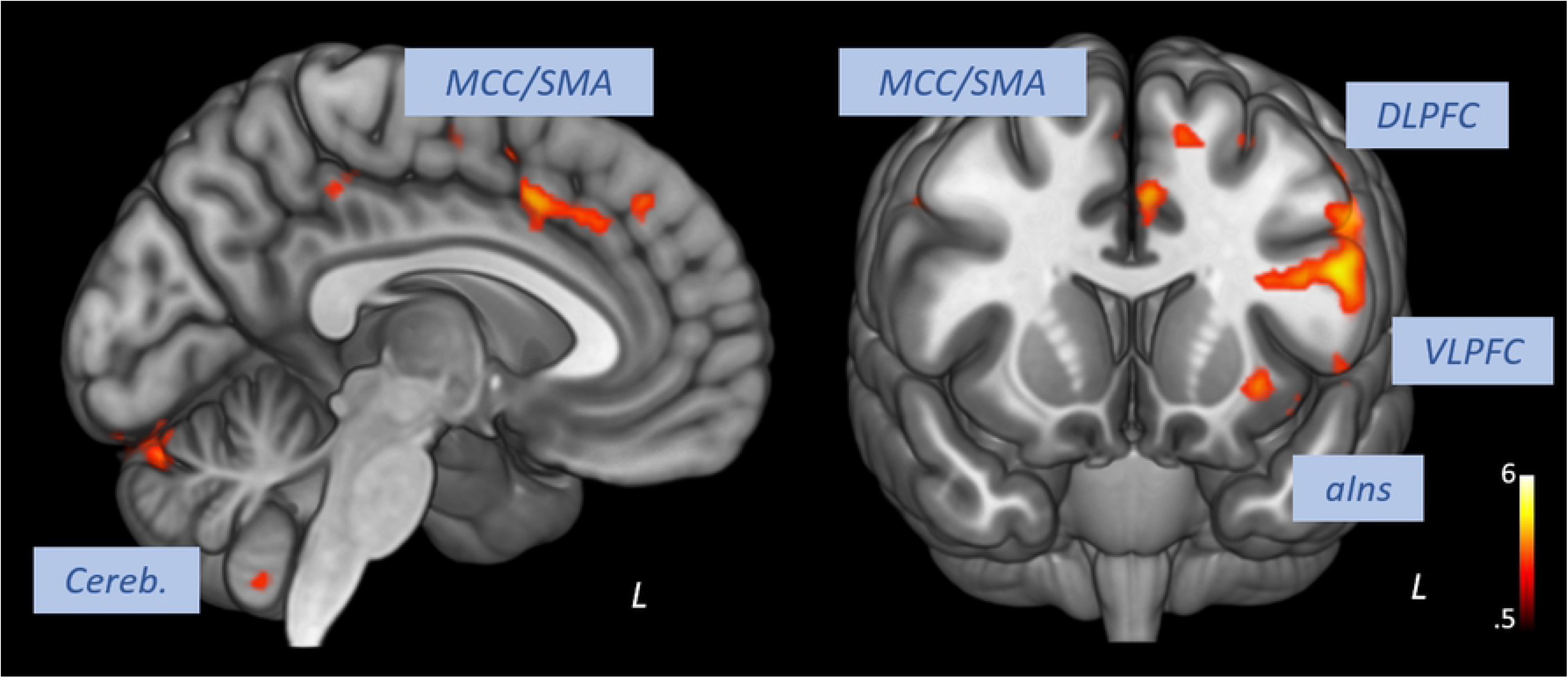
Correlation of heart rate variability changes with changes of functional connectivity of the prefrontal cortex. MCC: middle cingulate cortex, SMA: supplementary motor area, dACC: dorsal anterior cingulate cortex, DLPFC: dorsal lateral prefrontal cortex, aIns: anterior insula, Cereb: cerebellum (voxel-level: p<0.005 uncorr., cluster-level: k>10)

### Effect of biofeedback on heart rate variability

As depicted in Table 1, resting heart rate during functional scans decreased after the biofeedback intervention by 5.5 beats/min (8 %). Global (SDNN) and short term (RMSSD) HRV increased by 10.2 ms (19 %) and 10.1 ms (22 %) respectively after biofeedback, whereas the breathing rate did not change significantly. The control intervention had no significant effect on any of these parameters. There was a significant group effect on HR changes (T(22)=−2.59, p<0.05) and RMSSD changes (T(22)=2.53, p<0.05).

**Table 1.** Changes of heart rate variability and breathing rate from before (T1) to after the intervention (T2) in the biofeedback and control group. Parameters were assessed during resting fMRI. T1: before intervention, T2: after intervention, HR: heart rate, RMSSD: root mean square of successive heart beat interval differences, SDNN: standard deviation of heart beat intervals, BR: breathing rate

| Parameter | Biofeedback group |  |  |  | Control group |  |  |  |
| --- | --- | --- | --- | --- | --- | --- | --- | --- |
|  | T1 | T2 | T2-T1 | Significance | T1 | T2 | T2-T1 | Significance |
| HR [$\text{min}^{-1}$] | 70.2 ± 9.9 | 64.7 ± 8.7 | -5.5 ± 5.6 | p<0.05 | 70.2 ± 8.6 | 70.0 ± 6.6 | -0.2 ± 3.9 | n.s. |
| SDNN [ms] | 54.8 ± 15.4 | 65.0 ± 17.6 | 10.2 ± 12.1 | P<0.01 | 58.8 ± 26.4 | 60.7 ± 32.3 | 1.9 ± 10.4 | n.s. |
| RMSSD [ms] | 45 ± 18.3 | 55.1 ± 19.8 | 10.1 ± 7.1 | p<0.01 | 47.5 ± 27.5 | 49.6 ± 25.5 | 2.2 ± 8.1 | n.s. |
| BR [bpm] | 16.4 ± 4.2 | 15.1 ± 3 | -1.3 ± 3.9 | n.s. | 15.1 ± 3.2 | 15.3 ± 4.5 | 0.2 ± 2.6 | n.s. |

### Effect of biofeedback on functional connectivity of the prefrontal cortex

After the biofeedback intervention, we found increased connectivity between the VMPFC and the anterior and middle cingulate cortex, the supplementary motor area, dorsal and ventral lateral prefrontal regions, and the left anterior insula (see Figure 2A, Table S2). The group comparison revealed significantly higher increases of prefrontal connectivity to the anterior and middle cingulate cortex, left anterior insula, supplementary motor area, thalamus, dorsal and ventral lateral prefrontal regions in the biofeedback group when compared to the control group (Figure 2B, Table S3). Clusters in the pons (parabrachial nucleus) and ventral medulla (nucleus of the solitary tract) indicate stronger increases of functional connectivity of the VMPFC to vagal autonomic centers in the brainstem after the biofeedback intervention.

**Figure 2.**
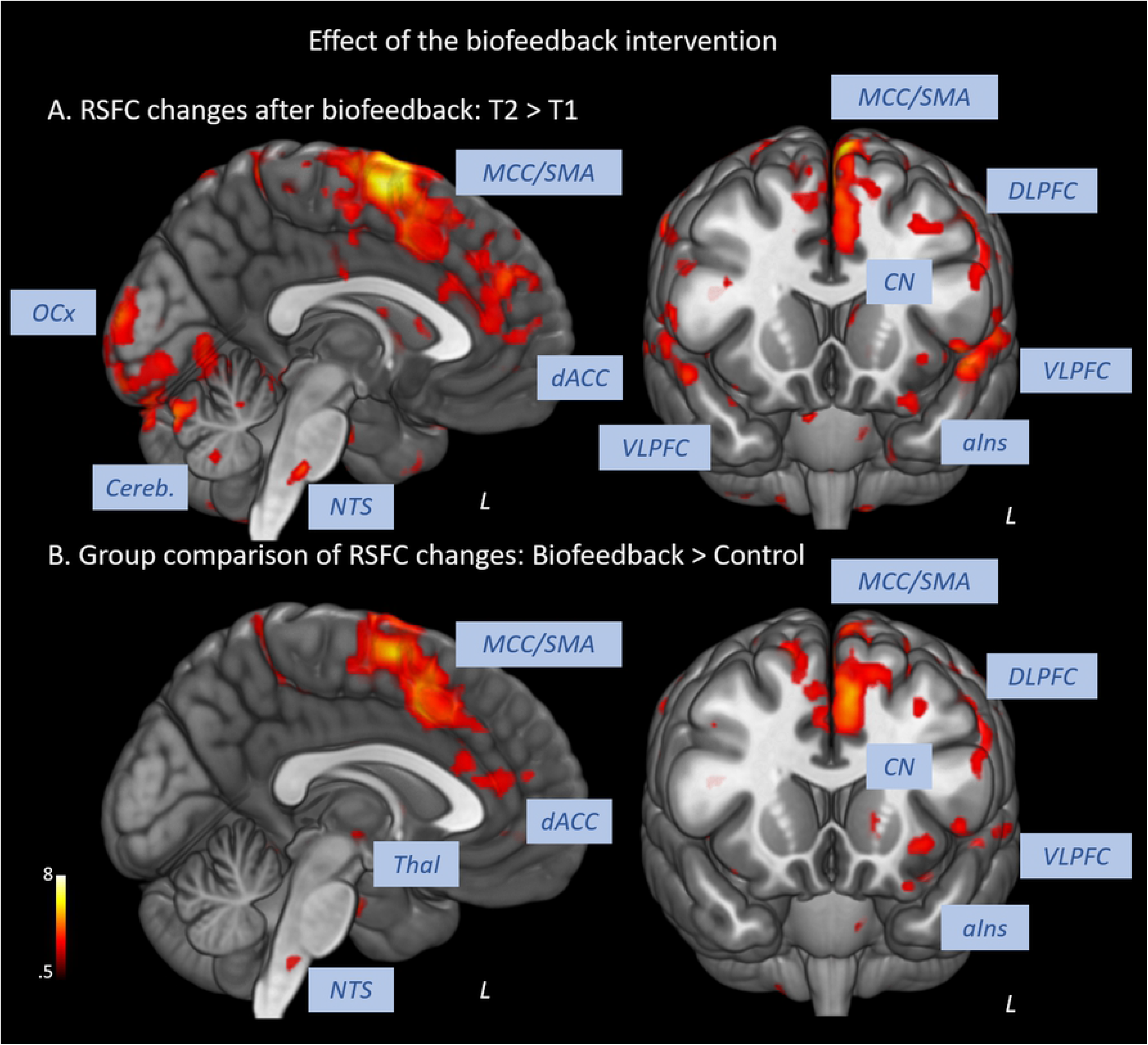
Effect of biofeedback on functional connectivity of the prefrontal cortex. A: Increases of functional connectivity from T1 to T2 within the biofeedback group. B: Significant increases of functional connectivity in the biofeedback group compared to control group. SMA/MCC: supplementary motor area / middle cingulate cortex, VLPFC: ventral lateral prefrontal cortex, DLPFC: dorsal lateral prefrontal cortex,Thal: thalamus, dACC: dorsal anterior cingulate cortex, aIns: anterior insula, OCx: occipital cortex, CN: caudate nucleus, Cereb: cerebellum, NTS: nucleus of the solitary tract (voxel-level: p<0.005 uncorr., cluster-level: k>10)

### Effect of biofeedback on functional network organization

Significantly greater positive functional connectivity was observed after the biofeedback intervention in a network of 33 nodes and 32 edges (Figure 3, p= 0.004) revealed by the NBS analysis. Nodes within this network were located in central autonomic regions, i.e. amygdala, ventromedial prefrontal cortex, anterior cingulate cortex, but also in visual, temporal and sensorimotor regions with a large number of intra-hemispheric functional connections.

**Figure 3.**
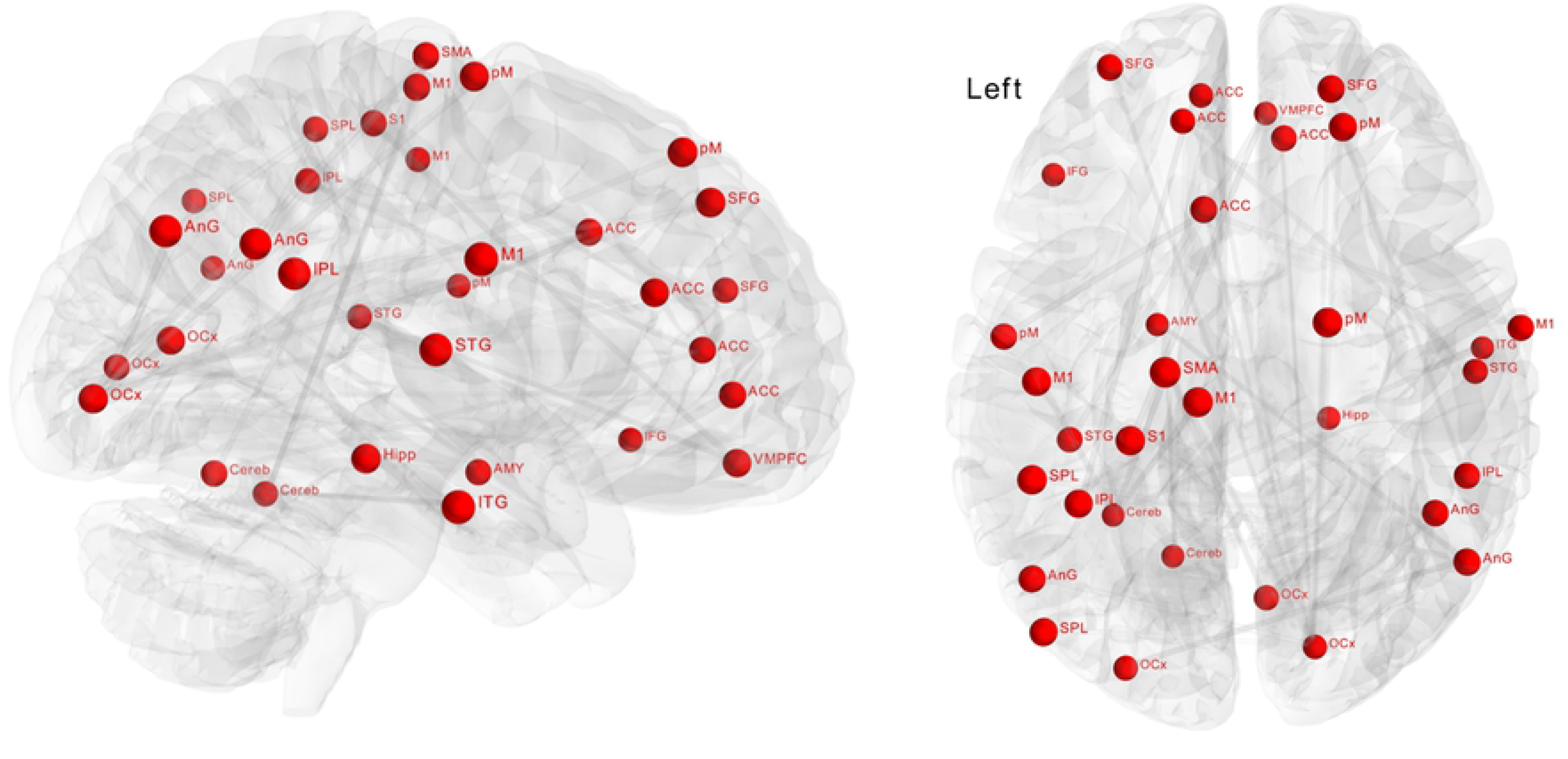
Effect of biofeedback on functional connectivity matrices using Network-based statistics (NBS). The depicted component shows nodes with significantly (p = 0.004) higher connectivity after biofeedback intervention. These connections formed a single connected network with 34 nodes and 32 edges.

## Discussion

Our results demonstrate increased HRV and decreased heart rates after HRV biofeedback training. Additionally, we found enhanced functional connectivity of the prefrontal cortex to a number of important cortical regions and autonomic centers in the brainstem. A wide functional network of brain regions seems to be affected by biofeedback intervention.

Biofeedback has been demonstrated to restore autonomic dysfunction in patients with cardiovascular disorders[15]. For example, in a group of patients with coronary artery disease, an HRV biofeedback intervention increased resting HRV, reduced blood pressure, and also decreased hostility behavior[32]. A number of studies also showed that HRV biofeedback reduces self-reported stress and anxiety with a large effect size that does not seem to be limited to a clinical level of anxiety[16]. Based on previous findings, linking high HRV to better response inhibition and emotion regulation[33–35], Mather & Thayer (2018) hypothesized that HRV biofeedback enhances interactions in emotion regulatory brain regions. Core brain regions that are involved in the experience and perception of emotions are the anterior insula, the amygdala, and the cingulate cortex[36]. The prefrontal cortex has regulatory control over these regions and determines the cognitive processing and interpretation of feelings[37,38].

Considering the CAN described by Benarroch (1993), a wide range of brain regions crucial for emotional processing are also involved in autonomic control. As emotional arousal is closely tied to autonomic responses, it is not surprising that their neural representations overlap[39,40]. Cognitive regulation via the prefrontal cortex can neutralize emotional affective experiences and decrease accompanying physiological arousal[41,42]. Our current results show that even at rest, regulatory control of the medial prefrontal cortex over limbic regions is closely related to HRV changes. We found that dynamic functional connectivity between the ventromedial prefrontal cortex and the anterior insula, the cingulate cortex and the amygdala correlated with the time course of HRV.

The HRV biofeedback intervention increased prefrontal functional connectivity, especially to the anterior insula, anterior cingulate cortex, thalamus, lateral prefrontal regions, and brainstem autonomic nuclei. According to the neurovisceral integration model, prefrontal control over the cingulate cortex and the insula reflects the highest level of a top-down regulatory chain involving the amygdala, hypothalamus, and the brainstem to modulate cardiac activity[43]. Sympatho-excitatory subcortical circuits are under tonic inhibitory control by the PFC[44]. For example, the amygdala, which has outputs to autonomic, endocrine, and other physiological regulatory systems and is activated during threat and uncertainty, is under tonic inhibitory control via GABAergic projections from the PFC[45,46]. Thus, in normal modern life, the sympatho-excitatory preparation for flight and fight is tonically inhibited. However, under conditions of uncertainty or threat, the PFC becomes hypoactive which is associated with disinhibition of sympatho-excitatory circuits that are essential for physical and mental responses. Similarly, it has been postulated that psychopathological states such as anxiety or depression are associated with prefrontal dysfunction leading to poor habituation to novel neutral stimuli or unbalanced threat information processing[47,48]. As a consequence, sympatho-excitatory circuits become disinhibited in these conditions leading to abnormal emotional processing as well as to an autonomic imbalance[49,50].

The enhancement of interactions between specific brain regions might underly the beneficial influence of HRV biofeedback on emotion regulation. Network-based statistics revealed that the connectivity in a wide network of regions was influenced by biofeedback with nodes located in the central autonomic network, but also in the visual and sensorimotor system.

How biofeedback of a peripheral autonomic signal influences functional brain organization is still unclear[14,17]. By adapting breathing in order to maximize heart rate oscillations, participants ‘exercise’ principle vagal reflexes, especially the baroreflex[51]. The baroreflex is one of the most powerful mechanisms of short-term heart rate modulation. Pressure sensors called baroreceptors detect changes of blood pressure and initiate adaptation of cardiovascular function. Immediate influences on heart rate are vagally mediated via autonomic centers in the brainstem[52].

By augmenting vagal afferent input, HRV biofeedback is thought to stimulate those cardiovagal brainstem nuclei similar to direct electrical stimulation[17]. Vagal nerve stimulation acts on the central autonomic network, and the limbic system by modulating vagal afferent activity[19]. The nucleus of the solitary tract (NTS) is the primary integration center of sensory information from the periphery, including discharge patterns of baroreceptors and lung stretch receptors, with projections to noradrenergic and serotonergic neuromodulatory systems[53,54]. Using fMRI, it has been demonstrated that vagal nerve stimulation increases activity of the NTS and enhances its functional connectivity to the midcingulate/SMA and anterior insula[55,56].

Our data support the involvement of brainstem autonomic nuclei, i.e. the NTS and the parabrachial nucleus, in mediating the effect of biofeedback on higher cortical regions. The medial parabrachial nucleus, especially the Pre-Bötzinger complex, acts as a respiratory pacemaker[57,58]. The lateral part is involved in coordination of the baroreflex[59]. However, spatial resolution of current functional scanning sequences is barely sufficient to identify BOLD signal changes in very small brainstem nuclei. Further studies of this area need to involve more sophisticated image processing methods[60].

## Conclusion

In conclusion, our data suggest that HRV biofeedback increases HRV and decreases heart rate. Changes of autonomic cardiac regulation are accompanied by enhanced functional connectivity of the prefrontal cortex to core regions of emotional and cognitive processing. We found evidence that the central effect is mediated by autonomic centers in the brainstem.

## Acknowledgements

AS is supported by the Interdisciplinary Center for Clinical Research Jena (MSP05-2019, www.uniklinikum-jena.de/izkf). The funders had no role in study design, data collection and analysis, decision to publish, or preparation of the manuscript.

## Supplementary material

Table S1 Temporal co-variation of prefrontal connectivity and heart rate variability (p<0.005 uncorr., cluster size k>10)

Table S2 Effect of biofeedback on functional connectivity of the prefrontal cortex: T2 >T1 (p<0.005 uncorr., cluster size k>10)

Table S3 Group comparison of functional connectivity changes: biofeedback > control group (p<0.005 uncorr., cluster size k>10)

